# Electro-mechanical transfection for non-viral primary immune cell engineering

**DOI:** 10.1101/2021.10.26.465897

**Authors:** Jessica M. Sido, James B. Hemphill, Rameech N. McCormack, Ross D. Beighley, Bethany F. Grant, Cullen R. Buie, Paulo A. Garcia

## Abstract

Non-viral approaches to transfection have emerged a viable option for gene transfer. Electro-mechanical transfection involving use of electric fields coupled with high fluid flow rates is a scalable strategy for cell therapy development and manufacturing. Unlike purely electric field-based or mechanical-based delivery methods, the combined effects result in delivery of genetic material at high efficiencies and low toxicity. This study focuses on delivery of reporter mRNA to show electro-mechanical transfection can be used successfully in human T cells. Rapid optimization of delivery to T cells was observed with efficiency over 90% and viability over 80%. Confirmation of optimized electro-mechanical transfection parameters was assessed in multiple use cases including a 50-fold scale up demonstration. Transcriptome and ontology analysis show that delivery, via electro-mechanical transfection, does not result in gene dysregulation. This study demonstrates that non-viral electro-mechanical transfection is an efficient and scalable method for cell and gene therapy engineering and development.

**One Sentence Summary:** This study demonstrates that non-viral electro-mechanical transfection is an efficient and scalable method for development of engineered cellular therapies.

## INTRODUCTION

Immunotherapy is currently at the cutting edge of basic scientific research and pharmaceutically driven clinical application. This trend is in part due to recent strides in targeted gene modification and expanded use of CRISPR/Cas complex editing for therapeutic development (*1*). Identification of genetic modifications of therapeutic interest requires screening thousands of genetic variants, which can include modification of an endogenous gene or insertion of an engineered gene (*2*). However, transfection steps for genetic modification are often limited to low throughput, inefficient technologies (*3*). Automated platforms for high efficiency transfection have the potential to reduce process costs substantially while increasing the number of successfully engineered cells. While a viral methodology can be applied to high-throughput automated systems, there are production limitations that extend timelines for research efforts: viral vectors have to be cloned and transfected into a viral production line, and viral particles must be purified prior to transduction (*4*). This process can take months, significantly affecting development timelines while simultaneously increasing the cost of drug discovery, and is not applicable for all cell types, specifically immune cell subsets resistant to viral infection.(*5*) As such there is an unmet need for a high-throughput automated gene transfer system that does not rely on viral delivery mechanisms.

Recent findings (2, 6, 7) suggest non-viral transfection represents the next step in the evolution of autologous and allogeneic cell therapy. Despite the advantages of separating cell therapy from viruses, non-viral transfection has lagged behind viral methodologies in clinical applications. One well-known form of non-viral transfection is electroporation, where a high energy electric field is applied to a static cell suspension (8, *9*). Successful transfection via classical static electroporation is dependent on the electric field strength each cell experiences (*10, 11*). However, electric fields that are too intense can result in irreversible cell membrane disruption, leading to cell death (*11*).

To improve the yield of electric field assisted transfection, the electrical energy applied to cells must be minimized. We posit that the electrical energy required to enable cell membrane permeability can be reduced by adding a mechanical component to the total energy applied to cells. This mechanical energy can be delivered via fluid flow, reducing the high energy electric fields needed for efficient delivery of genetic payloads (*2*). This paper introduces the *Flowfect^®^* transfection platform (Kytopen Corp; Cambridge, MA), an electro-mechanical method leveraging both high fluid flow rates and electrical energy for reversible pore formation.

Electro-mechanical cell transfection involves the use of electric fields coupled with mechanical stress associated with fluid flow rates that together enable cell permeation and delivery of exogenous material. This technique is distinct from electroporation, where an electric field is utilized to permeate cells typically with no flow or at low fluid flow rates that result in minimal stress, and purely mechanical cell transfection techniques, that utilize physical constrictions or novel flow geometries to induce pore formation. In the case of electro-mechanical cell transfection, pore formation is mediated by the combined effects of the electric field and mechanical energy input in the form of shear and normal stresses on the cell. One would expect that electro-mechanical cell transfection would depend upon the following parameters: the rootmean-square of the applied voltage, *V_RMS_*; medium conductivity, *σ*; average fluid velocity, *u*; the distance between electrodes, *l*; dynamic viscosity of the fluid, *μ*; the channel diameter, *d*; the cell diameter, *D*; and the fluid density, *ρ*. Applying the Buckingham Pi theorem for dimensional analysis (*12*), we obtain a set of four dimensionless parameters. The first two parameters 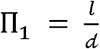 and 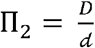, are dimensionless lengths that are scaled by the channel diameter. The third dimensionless group is the classic Reynolds Number, 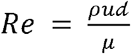, which is the ratio of inertial effects to viscous effects in the fluid flow. Electro-mechanical transfection typically occurs at moderate Re on the order 10^2^ and thus falls in the laminar flow regime (*13*). The final dimensionless group, 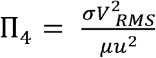, represents the ratio of electrical power applied to mechanical power imposed on the cell suspension.

We expect that the key physics of this process will be governed by these four dimensionless groups and combinations thereof. The existence and importance of Re and Π_4_ distinguish this transfection mechanism from both electroporation (*14, 15*) and purely mechanical based transfection methods under development (*16–21*). In electroporation the electric field and pulse conditions govern transfection efficiency, and the process typically occurs in static chambers with a quiescent fluid (*22, 23*). There are recent efforts involving flow-based electroporation in which the cell suspension moves with a finite velocity during the transfection process (*24, 25*). However, in these systems flow is used to deliver cells to the transfection zone, not to influence the transfection itself as is the case with electro-mechanical transfection. Mechanical based transfection methods, particularly those that employ higher flow rates (*20, 21*) would surely be dependent upon Re but as there is no applied electric field there is no need for Π_4_, described above. Therefore, despite its similarities to electroporation, we believe that electro-mechanical transfection is a novel technique to deliver exogenous material to cells.

## RESULTS

### Subhead 1: Application of electro-mechanical transfection

Electro-mechanical transfection was implemented with an automated liquid handler and a flow cell that integrates with the liquid handling system to enable delivery of electrical and mechanical energy to a cell suspension (Figure 1). These flow cells include a pipette tip architecture with a reservoir to enable aspiration/dispensing of cells and payload suspended in fluid buffer material. As the suspended cells and payload pass through the flow cell, with a defined flow rate, a precise electric field is delivered via contact with electrodes placed across the flow cell region. These cells are dispensed into a 96-well plate containing growth media and cultured for 24-hours. Analysis is then performed to determine output metrics via flow cytometer (gating example in Supplementary Figure 1).

**Fig. 1.**
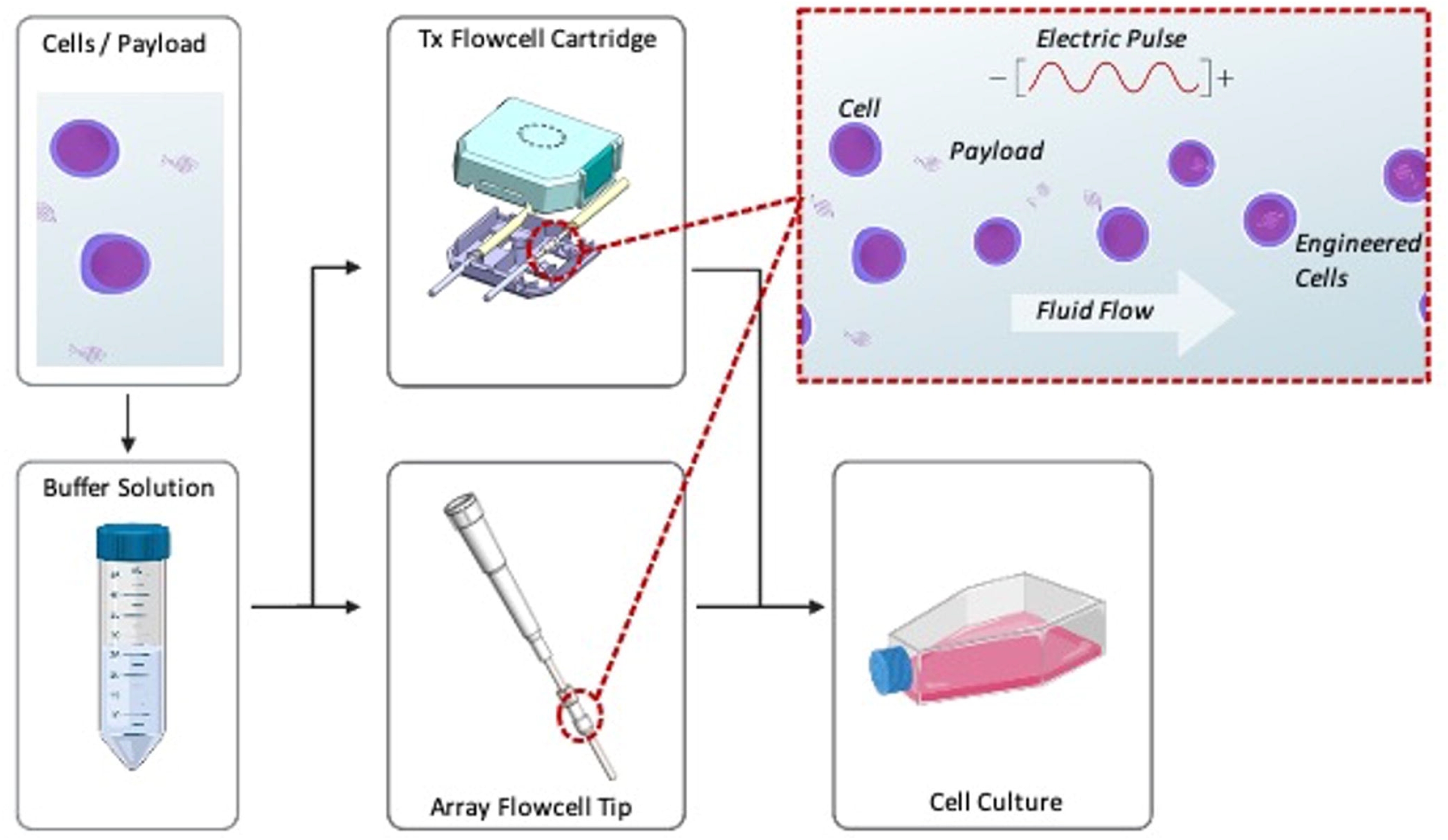
Schematic of our continuous flow non-viral Flowfect® platform. Cells & payload are suspended in proprietary buffer in the reservoir. As the cells & payload flow through the electro-mechanical zone they are exposed to both electric field energy and continuous fluid flow to induce transient cell membrane disruption and simultaneous delivery of genetic payloads into the cells. Transfected cells are then dispensed directly into growth media for cell recovery.

Effective use of electro-mechanical transfection requires determination of optimal conditions and parameters (e.g., flow rate and electric field) for the desired combination of cells and payload. To determine optimal conditions for application of the electro-mechanical transfection of an mRNA-based payload into primary human T cells (T cells), a plate-based matrix experiment was performed on a fully automated *Flowfect^®^* Array platform integrated with a commercial liquid handling system (PerkinElmer JANUS® G3 workstation, Waltham, MA). Utilizing this technology, up to 96 independently programmed combinations of electro-mechanical transfection parameters can be delivered. In this set of experiments an mRNA payload was delivered to 9-day expanded T cells 24-hours out of thaw. T cells were expanded using soluble anti-CD3/anti-CD28 antibodies and resuspended in transfection buffer supplied by Kytopen. The samples were recovered for culture and downstream analysis immediately after processing. Cell viability, live cell count, and transfection delivery efficiency (efficiency) were measured via flow cytometry (ThermoFisher Attune NxT); GFP (green fluorescent protein) reporter mRNA (GFP mRNA) was used to measure delivery efficiency to the T cells (Supplementary Figure 2). This set of experiments resulted in 11 conditions wherein transfected cells exhibit both high viability (over 70% live cells) and high efficiency (over 90% GFP^+^).

The most relevant cell transfection parameters, such as viability, efficiency, and cell yield are expected to depend upon the dimensionless parameters defined above including Π_1_, Π_2_, Re, Π_4_ and combinations thereof (Figure 2). Experience has shown that a fifth dimensionless group governs the dominant physics associated with cell transfection. We will denote this dimensionless group as 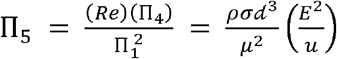. For a given cell type, in a particular transfection media, all terms outside of the parentheses can be considered a constant. Therefore, we expect cell transfection will typically vary most significantly with the applied electric field, *E*, and the average velocity in the channel, *u*. As shown in the data set below, we find that cell viability, efficiency, and cell yield, all appear to have a strong dependence on Π5. This data provides evidence that for a fixed channel geometry, media, and cell type, the value of Π_5_ is one of the principle factors determining cell transfection results. This factor, which couples the effects of both mechanical and electrical energy inputs, further indicates that electro-mechanical transfection is physically distinct from purely electrical or mechanical methods of cell transfection.

**Fig. 2.**
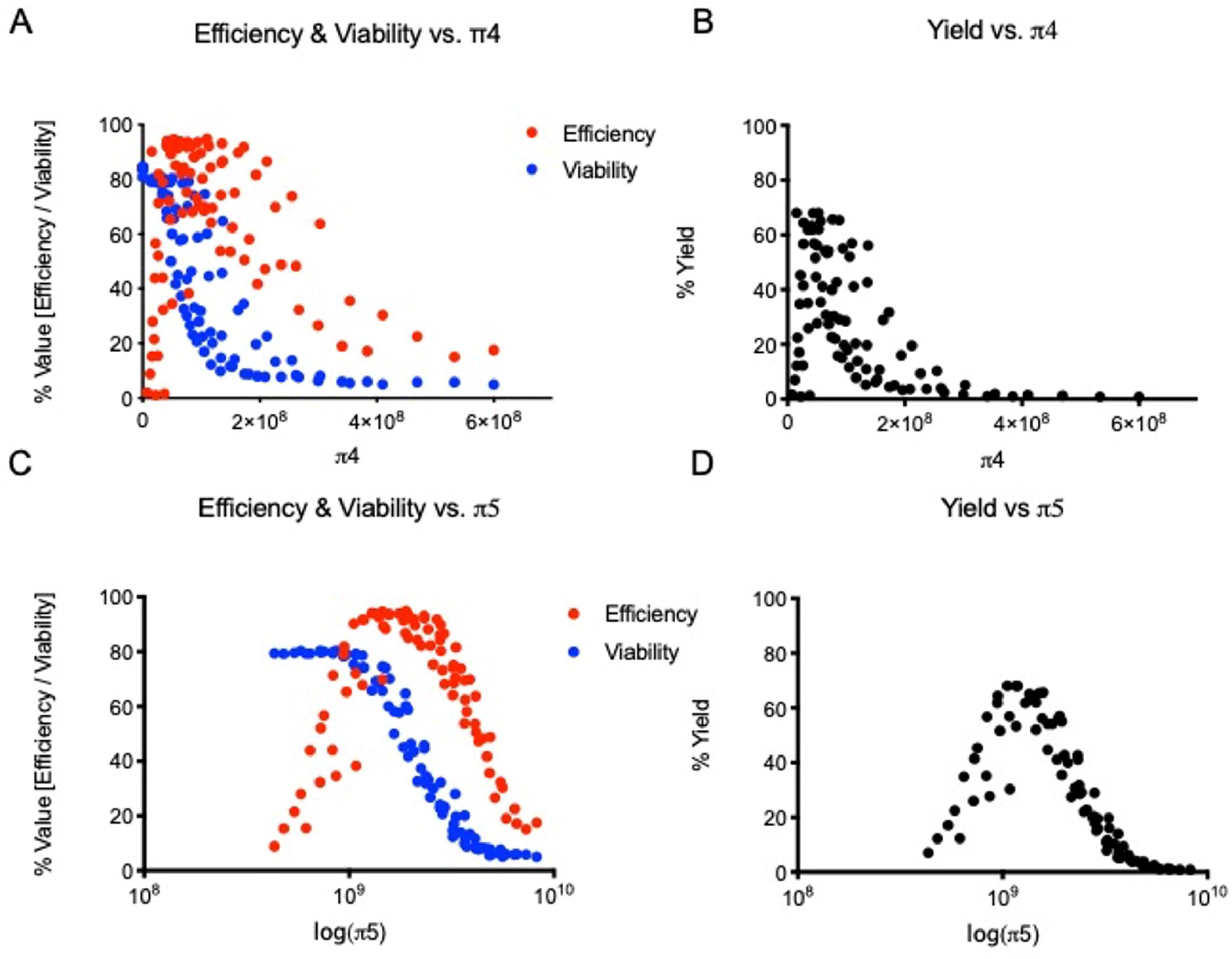
Parameters associated with electro-mechanical transfection correlated with outcome data. Expanded human T cells, transfected with a GFP reporter mRNA using our Flowfect® Array platform. Cultures were assessed for cell viability (7AAD negative), transfection efficiency, and percent yield (observed live GFP^+^ cells from 1e6 input cells) at 24 hours. The mechanism of action for Flowfect® transfection in GFP reporter mRNA delivery to expanded T cells is visualized here by plotting percent viability, efficiency, and yield against the electro-mechanical transfection specific parameter A & B) π4 and C & D) π5. All analysis was completed using the Thermo Fisher Attune NxT flow cytometer; n=89 depicted as individual data points

### Subhead 2: Transcriptional profiling

Transcriptome analysis was performed to further asses the effects of payload delivery into T cells using electro-mechanical transfection enabled via the *Flowfect^®^* technology (Figure 3). Commercially available non-viral electroporation systems were included for comparison metrics: The Neon™ transfection system from ThermoFisher (Neon) and the 4D Nucleofector™ from Lonza (4D Nucleofector). Each system was evaluated using 100 μL reactions containing 5M cells and device-specific proprietary programs and buffers. For each device, cells were processed without payload present and compared to a donor control that did not experience any processing. For this analysis, cells from two donors were processed in duplicate on each device. Significant (p<0.05) gene dysregulation, greater than 1-fold, at 6-hours after cell processing are shown in volcano plots as red (upregulated genes) or green (downregulated genes) in Fig. 3a-c (replicate data from the second donor is shown in Supplementary Figure 3). The *Flowfect^®^* system exhibited a nearly baseline gene expression profile at 6-hours, with only 2% of all genes dysregulated by the electro-mechanical transfection process (Figure 3d). The Neon exhibited a low dysregulation profile at 6-hours, with 6% of all genes dysregulated by this electroporation process (Figure 3d). The 4D Nucleofector exhibited significant dysregulation at 6-hours, with 47% of all genes dysregulated (Fig. 3d).

**Fig. 3.**
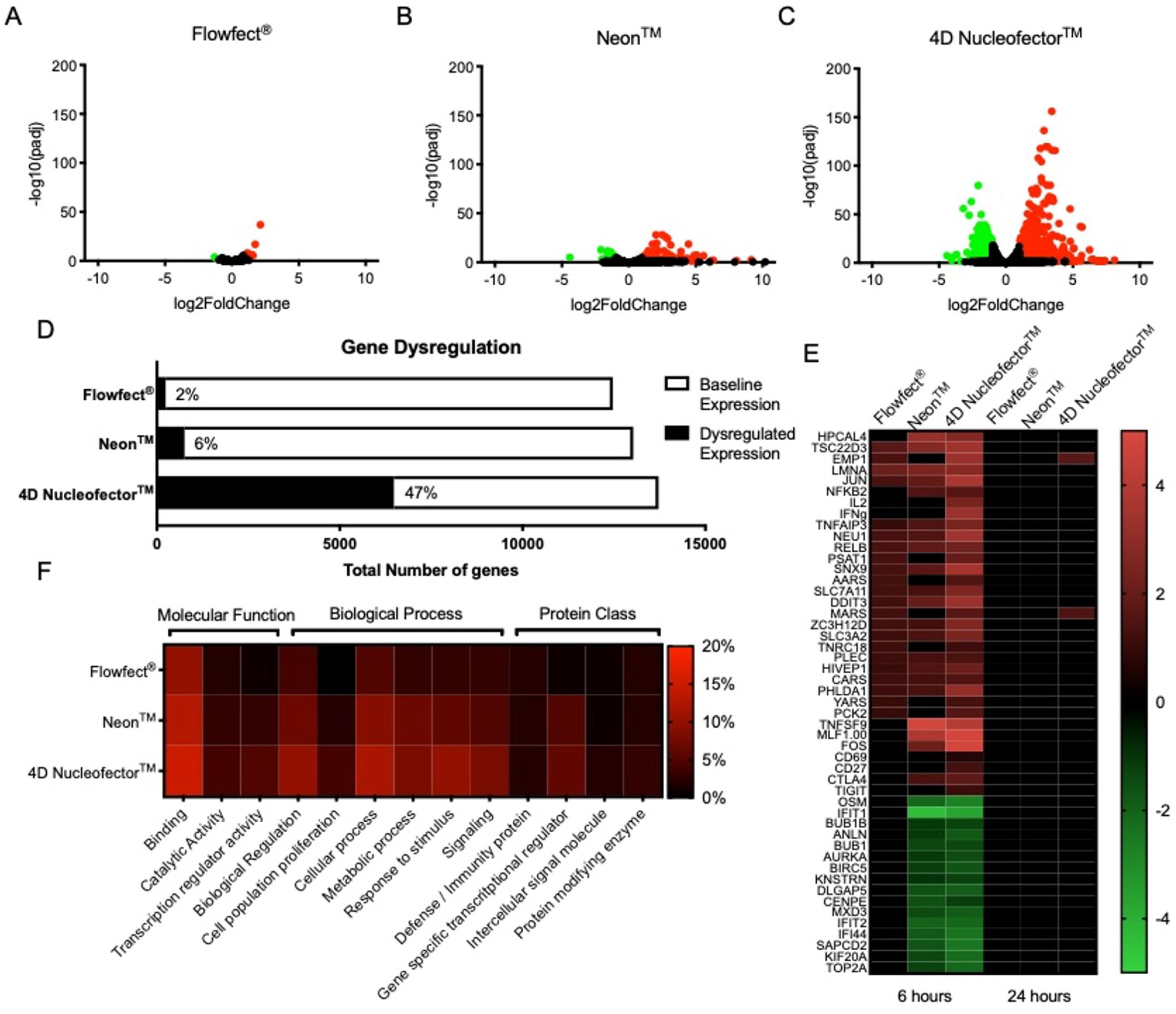
Electro-mechanical transfection can be used without a loss of cellular function. Expanded human T cells were processed without payload using the Flowfect® platform or commercially available electroporation systems (Neon™ and 4D Nucleofector™). Representative data is shown here at 6 or 24 hours post processing. A-C) Volcano plots showing significantly dysregulated genes (p<0.05) with greater than 1-fold change in expression at 6 hours. D) Graphical representation of genes exhibiting baseline versus dysregulated expression at 6 hours. E) Heatmap of selected up (red) and down (green) regulated genes at 6 and 24 hours. F) Heatmap of gene ontology focused on T cell function. Data shown for Donor # 1.

The functional capability of a cell product can directly impact the effectiveness of cells to drive desired immunological responses. For instance, it has been shown that edited cell differentiation and exhaustion can be linked to limited efficiency of T cell therapies (*26*). To better understand the impact of cell processing on function, upregulation of genes commonly associated with T cell function were assessed at 6-hour and 24-hour time points (Fig. 3e). We included proinflammatory cytokines (IFN*γ* and IL-2) and activation receptors (CD69 and CD27) to assess process driven activation of the cells (*27*). Exhaustion receptors (CTLA4 and TIGIT) were selected as indicators of process impact to cellular function downstream in treated cells (*28*). No upregulation of proinflammatory cytokines, activation receptors, or exhaustion markers were observed in electro-mechanically treated cells (Fig. 3e). Conversely, the Neon upregulated CTLA4 and the 4D Nucleofector upregulated proinflammatory cytokines, activation receptors, and exhaustion markers in treated cells (Fig. 3e). To further explore the impact to overall cell health and function, gene ontology was assessed in molecular function, biological process, and protein class using the Protein Analysis Through Evolutionary Relationships (PANTHER) classification system (*29*). At the 6-hour time point electro-mechanical transfection showed that 6% of the total dysregulation was associated with protein class, 13% was attributed to molecular function, and 18% of the total dysregulation was associated with biological processes (Fig. 3f). For the Neon, 10% of the total dysregulation was associated with protein class, 19% was attributed to molecular function, and 36% of the total dysregulation was associated with biological processes (Fig. 3f). 4D Nucleofection resulted in 13% of the total dysregulation being associated with protein class, 24% was attributed to molecular function, and 52% of the total dysregulation was associated with biological processes (Fig. 3f).

To correlate the transcriptome data with post-processing viability and delivery efficiency, cells from the same donors were transfected with payload using the same programs and conditions for the *Flowfect^®^*, Neon™, and 4D Nucleofector™ platforms (Supplementary Figure 4). The *Flowfect^®^* system exhibited high transfected cell viability near 80%, similar to the Neon, while the 4D Nucleofector exhibited low transfected cell viability near 45% (Supp. Fig. 4a). Regarding delivery, both the *Flowfect^®^* and Neon achieved high delivery efficiency near 90%, while the 4D Nucleofector resulted in moderate delivery efficiency near 50%. (Supp. Fig. 4b). Taken together with the transcriptome analysis we concluded that non-viral delivery efficiency is not tied to poor cell product health post processing. Moreover, electro-mechanical transfection compares favorably to existing electroporation devices in terms of all metrics, including gene dysregulation, viability, efficiency, and cell health metrics.

### Subhead 3: Delivery of multiple mRNAs to primary human T cells

We performed experiments to evaluate the capability of electro-mechanical transfection methods to deliver multiple payloads both in parallel (co-delivery via single treatment) and in series (staggered treatments 48-hours apart). The parallel condition was performed on the same day with a cell mix containing two mRNAs, including GFP and mCherry reporter mRNA, while the series treatments were performed two days apart with cell mixes containing a single mRNA at each time point, GFP then mCherry 48-hours later (Supplementary Figure 5). The viability of T cells 24-hours after treatment with electro-mechanical transfection was near 80% for both methods (Supp. Fig. 5a), demonstrating that neither parallel nor in series transfections were detrimental to cell health. Electro-mechanical transfection allows for repeat staggered transfections without significant loss in cell viability. Different expression profiles were observed for the two methods (Supp. Fig. 5b). Dual delivery efficiency for the parallel method was 94.2%; while efficiency was 82.3% when the transfection was performed in series (Supp. Fig. 5c). There was a clean 1:1 expression observed for co-delivery of mRNA in parallel with very few cells (1%) expressing only a single fluorophore (Supp. Fig. 5b). In contrast, with in series transfections 3.3% of the population were single positive for GFP and 11.3% of the population were single positive for mCherry (Supp. Fig. 5c).

### Subhead 4: Multiple T cell donors

Donor heterogeneity is a constant in all cell therapy manufacturing and development pipelines; necessitating that output metrics be assessed across multiple donors. For clinical manufacturing, source material including T cells from various donors can require re-characterization and comparability testing (*30*). Therefore, it is critical that cell therapy development demonstrate results achieved during optimization translates to T cells sourced from a variety of starting material (Figure 4). Cells from three different healthy PBMC donors (demographics listed in Supplementary Table 1) were isolated, expanded, and electro-mechanically transfected with GFP mRNA. All donors in this study met starting phenotypic and viability criteria. This experiment demonstrated consistent results with multiple donor cellular material with less than 10% change in viability from non-transfected controls (Fig. 4a), and GFP mRNA efficiency had low variability (over 84%) for all three donors (Fig. 4b).

**Fig. 4.**
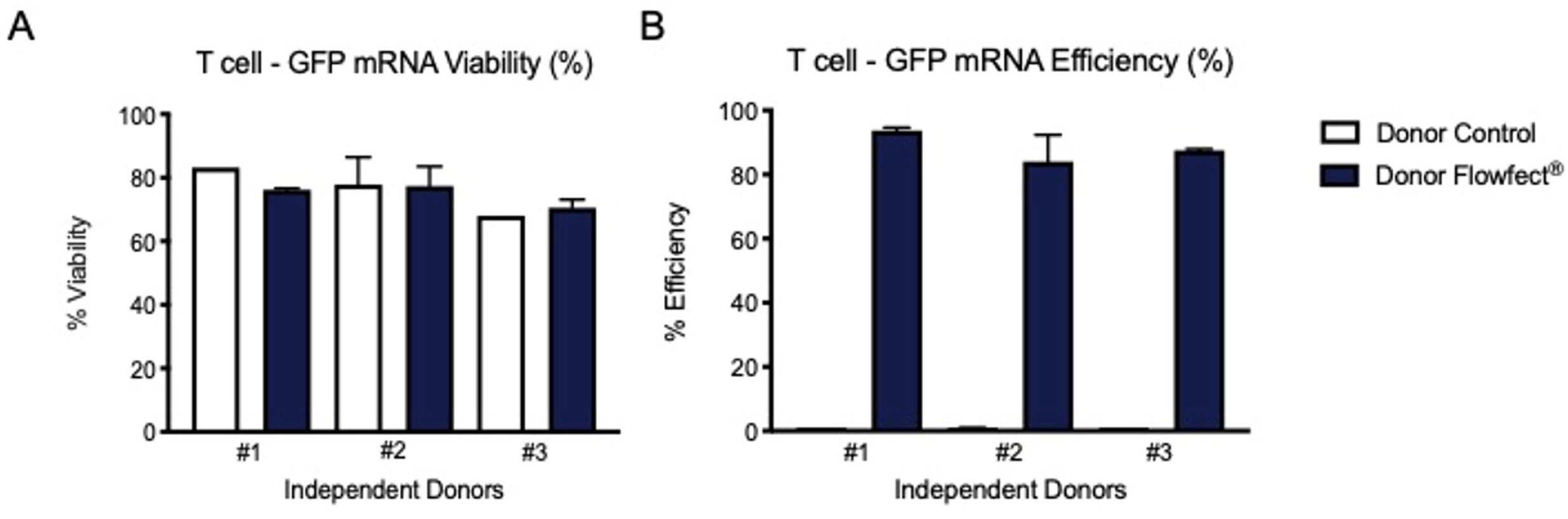
Flowfect® technology can be used across different donors. Expanded human T cells from three unique donors were transfected with a GFP reporter mRNA using the Flowfect® Array platform. Cultures were assessed for A) cell viability (7AAD negative) and B) transfection efficiency at 24 hours. Bar graphs are Mean ±SD with the following transfection sample sizes Donor #1 n=3 Donor #2 n=22 Donor #3 n=3.

### Subhead 5: Delivery of mRNA to naïve human T cells

Although non-activated naïve T cells are of interest in cell engineering this cell type is underrepresented in the field due to challenges in gene delivery and the inability to perform retroviral transduction without first activating the T cells (*31*). Here, the performance of electro-mechanical transfection with isolated naïve T cells (CD3^+^/CD4^+^/CD45RA^+^/CD45RO^-^) is evaluated using GFP mRNA delivery (Figure 5). The naïve T cells were then expanded with soluble anti-CD3/anti-CD28 activation reagents and monitored for 6-days after transfection. Growth rates of these cells after transfection were equivalent to non-processed control cells up to 6-days after activation (Fig. 5a) with no significant loss in viability (Fig. 5b). Additionally, the cells were stained for naïve T cell markers CD45RA and CD45RO (Fig. 5c). No change in phenotype for the transfected cells was observed with the cells retaining a naïve CD45RA^+^/CD45RO^-^ state. Viability of the transfected naïve T cells was equivalent to nontreated cells at 95.4% and 98.3% respectively (Fig. 5d). Efficiency was observed at 96.7% (Fig. 5e), corresponding to a high total yield of 74.9% of input cells (Fig. 5f).

**Fig. 5.**
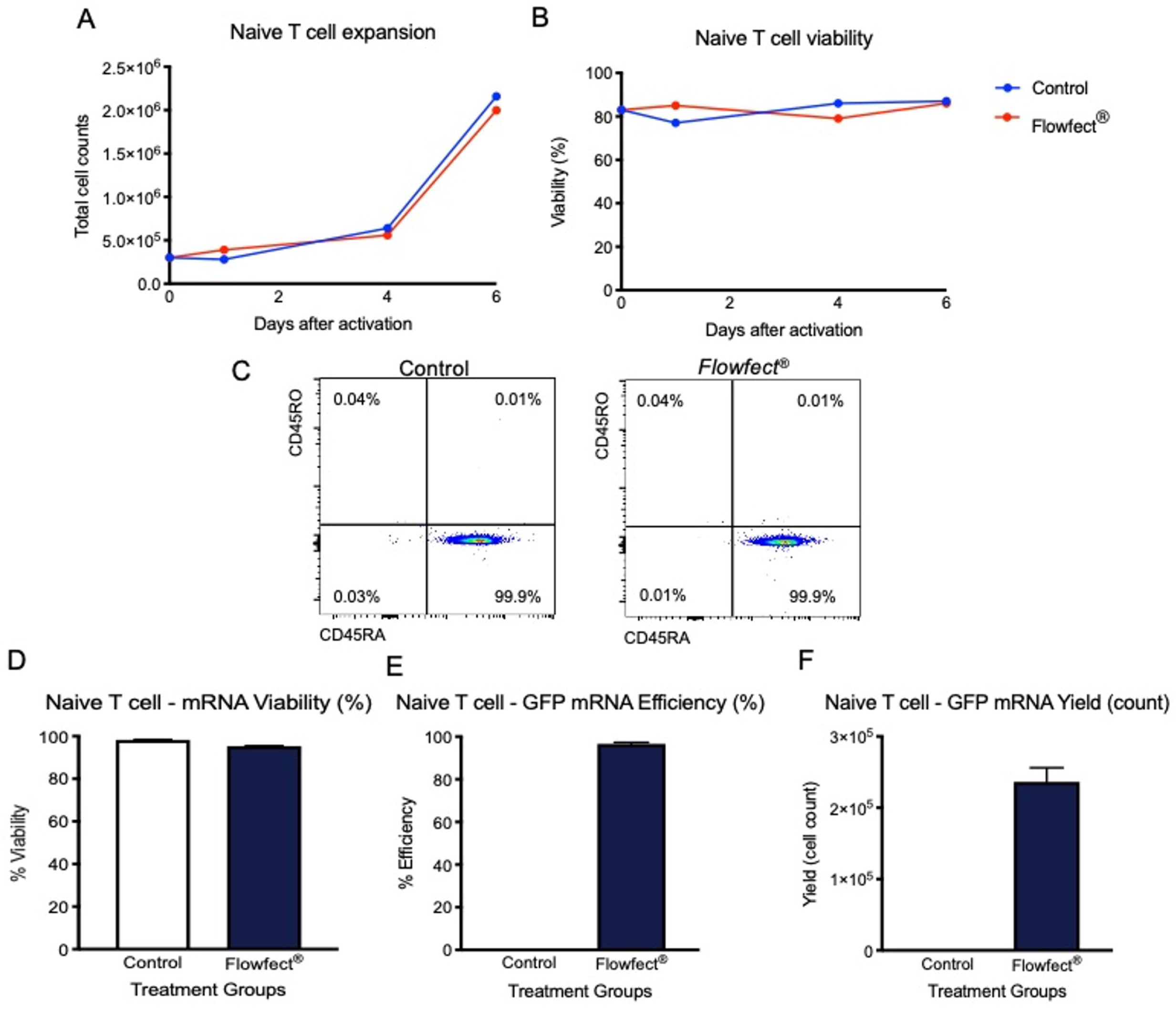
Flowfect® technology can be used to transfect naive T cells. Naïve T cells were transfected with a GFP reporter mRNA using our Flowfect® Array platform. Cultures were assessed for A) expansion capability and B) cell viability (Trypan blue exclusion) measured out to 6 days after transfection. The naïve T cells were also C) stained for expression of lineage markers CD45RA and CD45RO. Lastly, the D) cell viability (7AAD negative), E) transfection efficiency, and F) live GFP^+^ cell count (yield) from 0.5M cell input were measured at 24 hours after transfection. Bar graphs are representative of n=2 transfections Mean±SD.

### Subhead 6: Manufacturing volume scale-up

It has generally been accepted that electroporation requires additional optimization in the process of scaling up from research to manufacturing volumes, ascribed to changing geometries of both electrodes and cuvettes (*11*). To combat this issue, the field of electroporation-based transfection has seen the advent of workarounds including microfluidics, batch-based automation, and nanostructures (*32, 16*). However, these solutions have been unable to meet the need for high-throughput development and large volume manufacturing requirements in the evolving cell and gene therapy industry. Electro-mechanical transfection scales with time, therefore processing larger volumes only requires operating for a proportional length of time. To this end electromechanical transfection can process up to 100 mL of fluid (10-100B cells, depending on cell concentration) in roughly 3 minutes, from input to output bag, via a peristaltic pump (Masterflex® L/S). The transfection parameters identified during optimization on the small-scale system (*Flowfect^®^* Array) directly apply to the larger volume system (*Flowfect^®^* Tx) because these systems utilize the exact same electro-mechanical transfection apparatus (Fig. 6a). In a 50-fold scale up demonstration (5 mL) we transfected 50M T cells at a density of 10e6/mL with 1 mg of GFP mRNA (Fig. 6 b & c). The results show no significant loss in cell viability 24-hours after electro-mechanical transfection, with viabilities of 73.5% and 71.0% in small and large platforms respectively (Fig. 6b). The observed efficiency was also similar 24-hours after electromechanical transfection, 94.3% and 92.2% in small and large platforms respectively (Fig. 6c). Thus, electro-mechanical transfection can easily scale up for clinically relevant processing volumes.

**Fig. 6.**
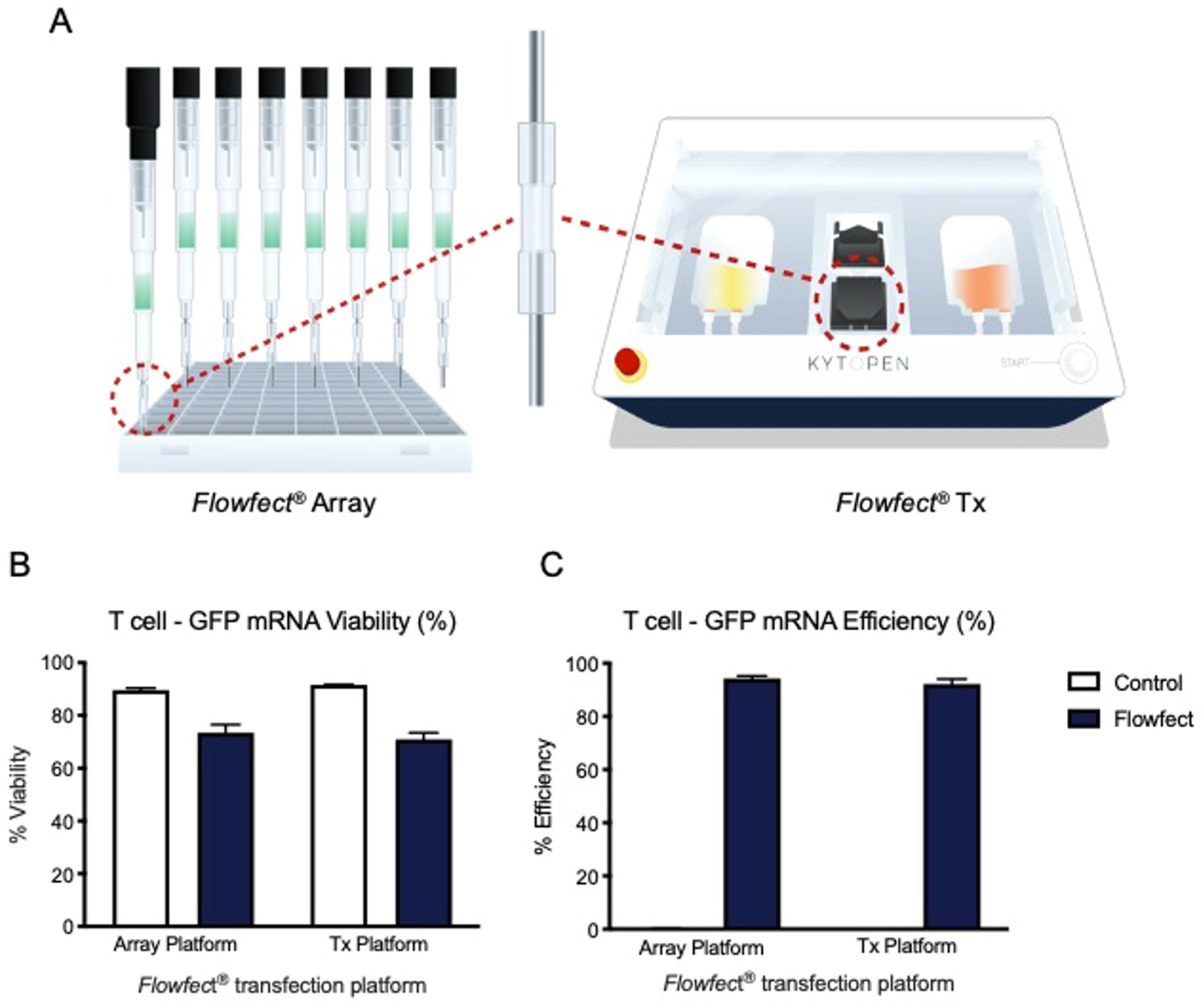
Flowfect® technology directly translates from small-scale research transfections to large-scale cell manufacturing transfections. **A)** Schematic showing how the electro-mechanical flowcell is shared between the Flowfect® Array and Flowfect® Tx, allowing for direct translation from one device to the other. Expanded human T cells were transfected with GFP mRNA reporter payload using our Flowfect® Array platform (small scale 100 μL transfections) and our Flowfect® Tx (large scale 5 mL transfections). The cells were assessed for B) cell viability (7AAD negative) and C) transfection efficiency at 24 hours. Bar graphs are Mean±SD Array n=6 Tx n=2.

## DISCUSSION

Non-viral transfection is an attractive method of engineering cells. This work presents a new transfection technology utilizing electrical energy with continuous flow that demonstrates several advantages over other non-viral transfection methodologies. Dimensional analysis reveals that electro-mechanical transfection is optimized by balancing effects of fluid flow and electric fields, distinguishing this technology from previous methods employing electric fields only. The physical model presented illuminates the critical parameters driving the effect of this technology, mainly a single dimensionless group containing the ratio of the electric field squared and the channel velocity. Further, this work provides an effective method for rapid optimization of the key parameters for effective electro-mechanical transfection.

The transcriptome analysis results show that high efficiency can be decoupled from significant gene dysregulation. Electro-mechanical transfection exhibited less than a 5% shift from baseline 6-hours after processing while tested commercially available electroporation devices exhibited greater than 5% shift from baseline in total gene dysregulation. Of the dysregulation induced by non-viral processing only 13% could be attributed to altered molecular function in the electromechanical device (*Flowfect^®^*) while electroporation devices (Neon™ and 4D Nucleofector™) induced between 19-24% dysregulation attributed to molecular function. This functional dysregulation was highlighted when markers of T cell exhaustion were found to be upregulated 6-hours after processing with electroporation devices but at baseline levels after processing with the electro-mechanical device. Analysis at 24-hours for efficiency showed that reduced gene dysregulation observed after electro-mechanical processing resulted in output metrics that were not significantly different from the commercially available Neon™ electroporation device, 89.2% and 89.4% respectively. Post transfection viability of greater than 75% and efficiency greater than 80% was observed in multiple use cases for electro-mechanical transfection. These findings were confirmed in multiple PBMC donors with no significant difference in efficiency, ranging from 84.0% - 93.7% for all three donors. The observed high efficiencies did not result in altered cell state as indicated in this study by maintenance of lineage specific naïve cell marker expression. Specifically, that high viability and efficiency, both above 95%, could be maintained in naïve CD4^+^ T cells with retention of naïve marker expression, 100% CD45RA^+^/CD45RO^-^. Moreover, results from the 50-fold scale-up transfection (*Flowfect^®^* Tx) resulted in ≤2.5% change in both viability and efficiency compared to small-scale results (*Flowfect^®^* Array).

While cell therapy promises significant efficacy advantages over standard of care therapeutics, and represents hope for individuals resistant to current options, change takes time. Only one FDA approved cell therapy solution (KYMRIAH, from Novartis) was released before 2020. As more cell and gene therapy clinical trials complete phase III testing only four names have been added to this exclusive list (Yescarta and Tecartus from Kite Pharma as well as Breyanzi and Abecma from Bristol Myers Squibb). This is in part due to the extended timelines and high costs of cell manufacturing via viral vectors. As the scientific community experiences the reduced timelines of moving from preclinical research to clinical testing with non-viral solutions such as electroporation (NCT03608618) and mechanical transfection (NCT04084951), the last obstacle will be scalability. The findings outlined in this study show that electro-mechanical transfection can compare favorably with, and sometimes exceed, the benchmarks of other non-viral options while decreasing the negative impact to throughput and cell product viability. The goal of this study was to define the capabilities of electro-mechanical transfection as a non-viral solution for human primary cell editing by evaluating delivery of mRNA into T cells. Further, studies advancing toward preclinical testing to characterize the health and function of electro-mechanically transfected cells in murine models are already underway. We seek to demonstrate functionality of the cell product, including in vivo efficacy and safety, prior to clinical manufacturing of engineered cells for human administration. Additionally, other clinically relevant exogenous materials and cell types must be evaluated in future work.

Together the data presented suggests that cell engineering using non-viral electro-mechanical transfection methods are distinct from classical static electroporation and represent a meaningful alternative to existing transfection methods. Electro-mechanical transfection can be leveraged with high throughput automation for discovery or process development, while also easily scaling up for clinical manufacturing. This ability for parallelization and scale up, while maintaining cell health and high cell yield, make electro-mechanical transfection an attractive new solution for cell therapy development and clinical manufacturing.

## MATERIALS AND METHODS

### Study design

Previous studies had indicated that the microfluidic electroporation concept underlying the electro-mechanical technology, which was initially explored in bacteria (*33–35*), could be applied to non-viral transfection of mammalian cells. However, the functional implications of electro-mechanical transfection and whether the resulting cell product could be utilized was unknown. To this end, our primary objective was to rigorously evaluate the mechanism of electromechanical transfection and the impact of high delivery efficacy in primary T cells. We selected primary T cells as a high impact cell type based on the broad applications possible in the immunotherapy space for T cell-based therapeutics (*36, 37*). The use of the GFP reporter system for delivery was based on the growing use of mRNA to avoid cell toxicity and possible longterm implications associated with DNA integration (*38*).

### T cell culture and expansion

Human peripheral blood cells (PBMCs) were purchased from STEMCELL Technologies (#70025). 100M PBMCs were thawed in 100 mL X-VIVO™ 10 media from Lonza (#04-380Q) with recombinant human IL-2 protein from R&D Systems (#202-IL). After a 24 hr culture out of thaw the PBMCs were activated with ImmunoCult™ human CD3/CD28 T cell activator reagent from STEMCELL Technologies (#10971) for 3 days according to the manufacturer’s protocol. On day 4 the cells were pelleted (500 x g, 5 min) and transferred to a G-Rex100® from Wilson Wolf (#80500) with 500 mL fresh X-VIVO™10 media and recombinant human IL-2. Fresh recombinant human IL-2 was then added every 3 days for up to 12 total days in G-Rex culture. Cells were then frozen into aliquots for future use with Bambanker cell freezing medium from Bulldog Bio (#BB01). Post expansion thawed aliquots of T cells were grown at 1e6/mL density in RPMI 1640 media from Thermo Fisher (#11875119) with 10% fetal bovine serum (FBS) from Sigma Aldrich (#F-4135), penicillin-streptomycin solution from Corning (#30-002-CI) and recombinant IL-2. Naïve primary human T cells (CD3^+^/CD4^+^/CD45RA^+^) were also sourced from STEMCELL Technologies (#70029) and cultured in RPMI with 10% FBS and IL-2 as described above. Cells were cultured at 37°C with 5% CO_2_ in a standard cell incubator. Cell viability and size were monitored during cell culture using Countess™ II (ThermoFisher).

### *Flowfect^®^* transfections

Transfections were performed with commercially sourced mRNAs encoding either GFP (#L7601) or mCherry (#L7203) from TriLink Biotechnologies. T cells were counted, pelleted (500 x g, 5 min), and resuspended in Kytopen’s proprietary *Flowfect^®^* transfection buffer at densities of 10-50e6/mL. Payload was added at a fixed maximum of 10% volume and the cell:payload solution was mixed via pipetting.

Processing with the *Flowfect^®^* Array were performed on a PerkinElmer JANUS® G3 BioTx Pro Plus Workstation with an 8-tip Varispan head. The cell solutions were transferred to a 4°C cooled mixing plate in a 96 well plate and aspirated through the *Flowfect^®^* Array tips above the microfluidic channel. The solutions were then dispensed through the same tips at constant flow rates while specific electric fields were applied to these tips through an electric delivery manifold that is a component of the *Flowfect^®^* System. The cells are delivered directly into cell culture media for recovery in a 96 deep well plate.

Processing with the *Flowfect^®^* Tx were performed with a prototype device developed by Kytopen in which fluid flows through a proprietary *Flowfect^®^* channel placed between input and output bags connected by tubing, with fluid transfer controlled using a Masterflex® L/S peristaltic pump and PharMed® BPT tubing (L/S 13: #06508-13) from Cole-Palmer. The cell:payload solutions were transferred to an input vessel and then pumped through the channel at constant flow rates while specific electric fields were applied. The cells were then immediately transferred into cell culture media for recovery in an output vessel.

Final density in recovery solution for both platforms was 1e6/mL, containing 10% *Flowfect^®^* transfection buffer and 90% cell culture media. Cells were cultured at 37°C with 5% CO_2_ in a standard cell incubator.

### Neon™ transfections

The ThermoFisher Neon™ transfection system was used according to manufacturer’s instructions. Briefly, 5M day 9 expanded T cells were resuspended in T buffer and loaded into the 100 μL Neon™ Pipette tip. The protocol ran was from the Neon™ T cell microporation protocol (2100V 1 pulse 20 ms). Processed cells were then transferred into a tissue culture vessel.

### 4D Nucleofector™ transfections

The Lonza 4D Nucleofector™ system was used according to manufacturer’s instructions. Briefly, 5M day 9 expanded T cells were resuspended in freshly prepared human T cell nucleofection solution and were loaded into the 100 μL Lonza certified cuvette. The protocol ran was from the Amaxa™ 4D Nucleofector™ protocol for unstimulated human T cells (EO115). Processed cells were then transferred into a tissue culture vessel.

### Flow cytometry analysis

A Thermo Fisher Attune Nxt flow cytometer was used for assessment of viability and efficiency metrics. 200 μL of cultured cells were pelleted (500 x g, 5 min) and resuspended in Dulbecco’s phosphate-buffered saline (DPBS) from Fisher Scientific (#14190250) with 7-AAD viability solution from eBiosciences (#00-6993-50). The cells were then analyzed on a volumetric read using the Attune Nxt autosampler. Total cells were gated in the forward scatter (FSC) and side scatter (SSC) dot plots. Viable cells (7-AAD^-^) were then gated to determine delivery efficiency via expression of the fluorescent reporters. This gating strategy is shown in Supplementary Figure 1. Total cell counts and yields were calculated from an applied dilution factor based on total volume of the cell culture (7.5X per 1 mL). Naïve T cell marker staining was performed with APC/Cy7 mouse anti-human CD45RA (#304128) and BV510 mouse anti-human CD45RO (#304232) antibodies from Biolegend.

### Transcriptome analysis

Whole cell pellets were collected from cell cultures at 6-hours and 24-hours post processing and stored at −80°C. All extraction of RNA, cDNA synthesis, next generation sequencing, and preliminary raw data normalized to controls was completed by GENEWIZ. Normalized data was then analyzed (Excel – Microsoft) and graphed (GraphPad – Prism 8) in house. Protein Analysis Through Evolutionary Relationships (PANTHER) classification system was used for gene ontology analysis.

### Statistical analysis paragraph

Data were statistically analyzed using JMP and Prism – GraphPad statistical software. Bar graphs are depicted as means. Error bars indicate SD unless otherwise indicated. Gene dysregulation was calculated using log2 fold change and significance calculated as -log10(padj).

## Supporting information

Fig. S1) Gating strategy for determining total cell counts and viability

Fig. S4) Electro-mechanical transfection compares favorably to commercially available non-viral transfection output metrics

Table S1) Unique donor demographics supporting data

## NOTES

## Acknowledgments

We thank PerkinElmer for providing an initial loan of the JANUS G3 BioTx Pro workstation and providing kits, consumables, and engineering support. The schematic in Figure 1 was created with BioRender.com. The *Flowfect^®^* schematics were created by The Like Minded (Marlow, UK). All work in this manuscript was completed at The Engine Accelerator (Cambridge, MA).

## Funding

National Science Foundation (NSF), Phase I SBIR Grant No. 1747096 (PG, RM, RB) National Science Foundation (NSF), Phase II SBIR Grant. No 1853194 (PG, RM, JH, JS, RB, BG)

MassVentures SBIR TARgeted Technologies (START) program Stage I

MassVentures SBIR TARgeted Technologies (START) program Stage II

## Author contributions

Conceptualization: JS, JH, CB, PG

Methodology: JS, JH, RM, RB, BG, CB, PG

Investigation: JS, JH, RM, RB

Analysis/Visualization: JS, JH, RM, RB, BG, CB

Funding acquisition: BG, CB, PG

Project administration: BG

Supervision: BG, PG

Writing – original draft: JS, JH, CB

Writing – review & editing: JS, JH, BG, CB, PG

