## Supplementary figures and images for "Electro-mechanical transfection for non-viral primary immune cell engineering"

### Fig. S1) Gating strategy for determining total cell counts and viability

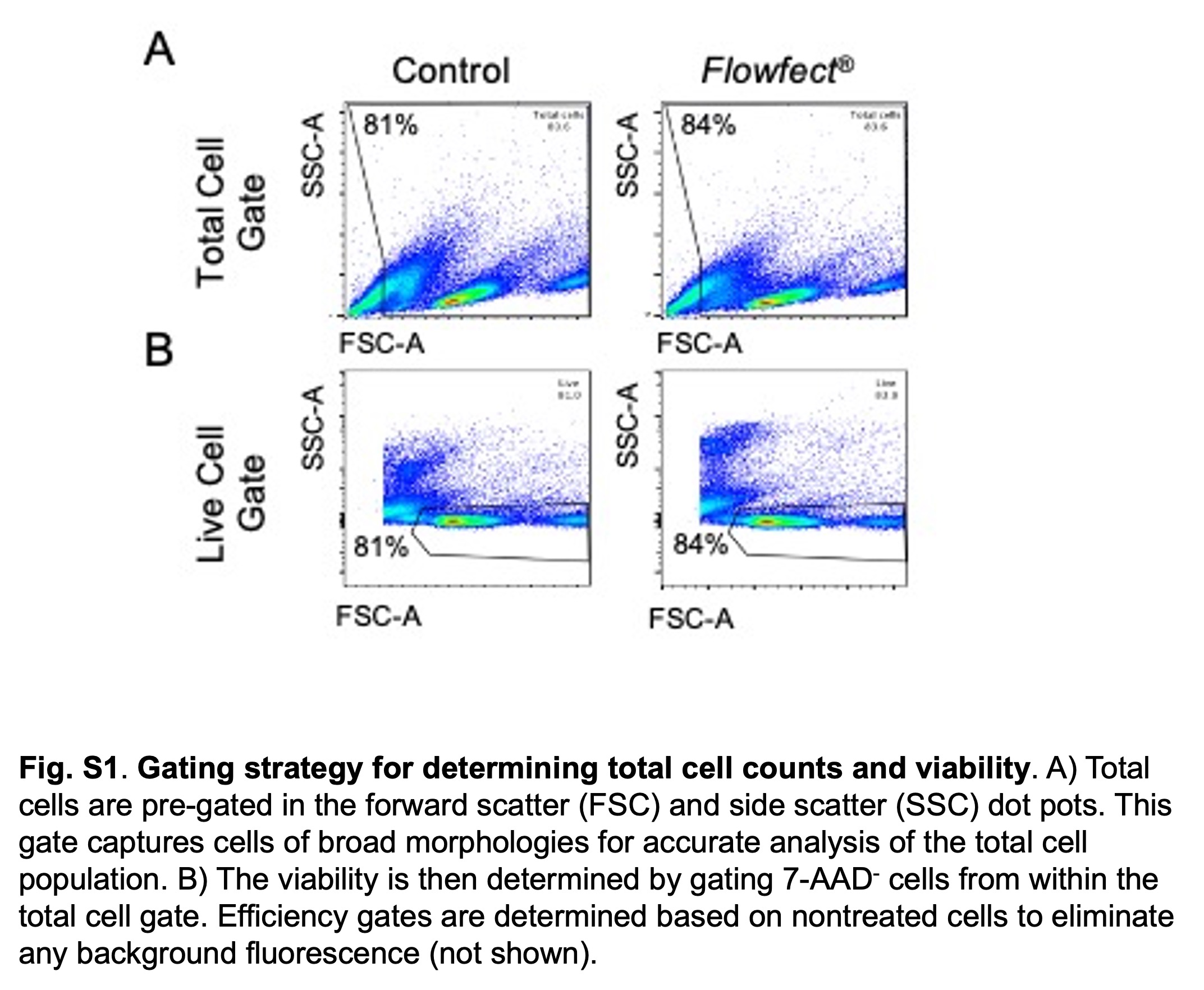

### Fig. S4) Electro-mechanical transfection compares favorably to commercially available non-viral transfection output metrics

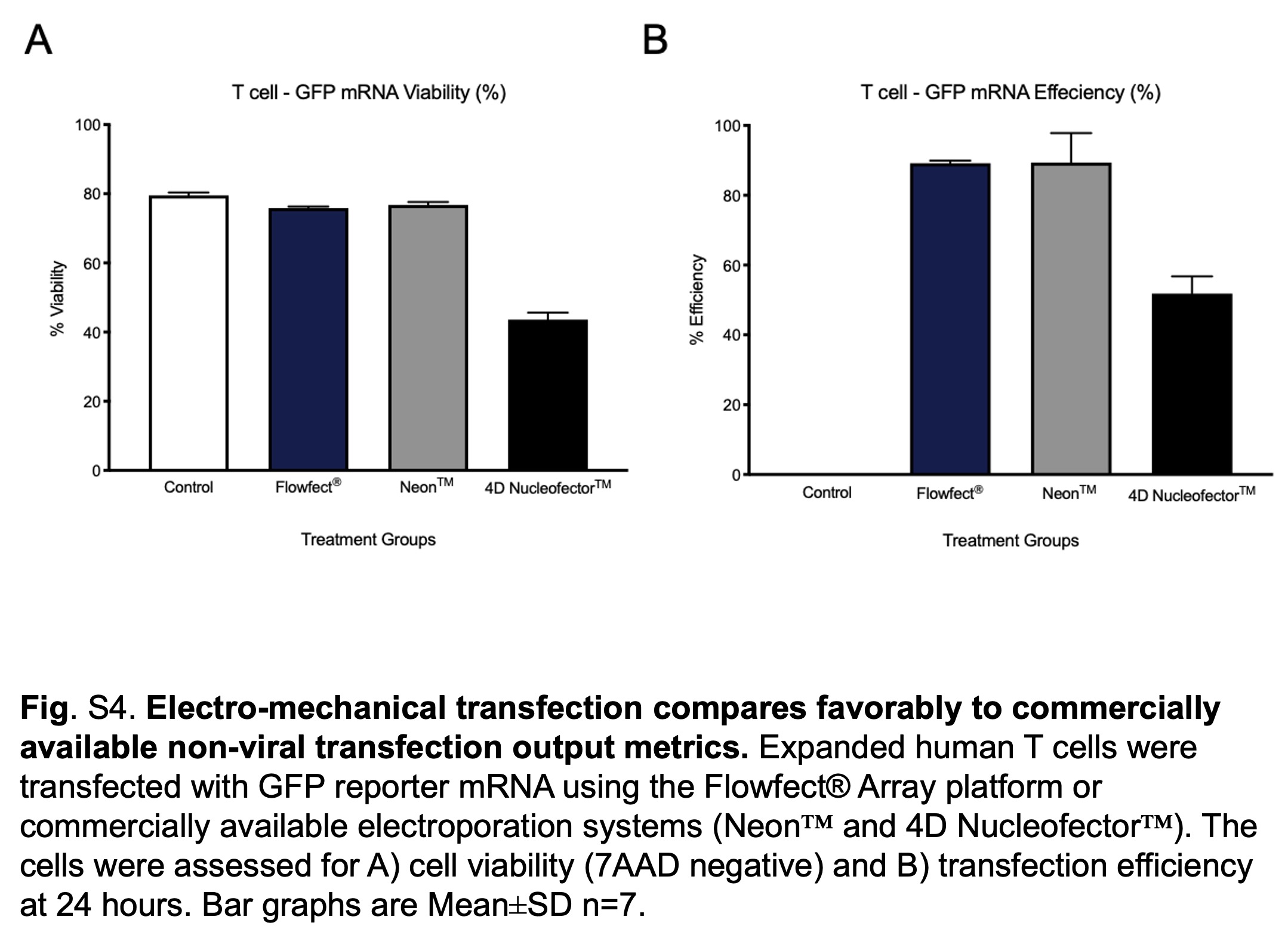

### Supplemental Data 1

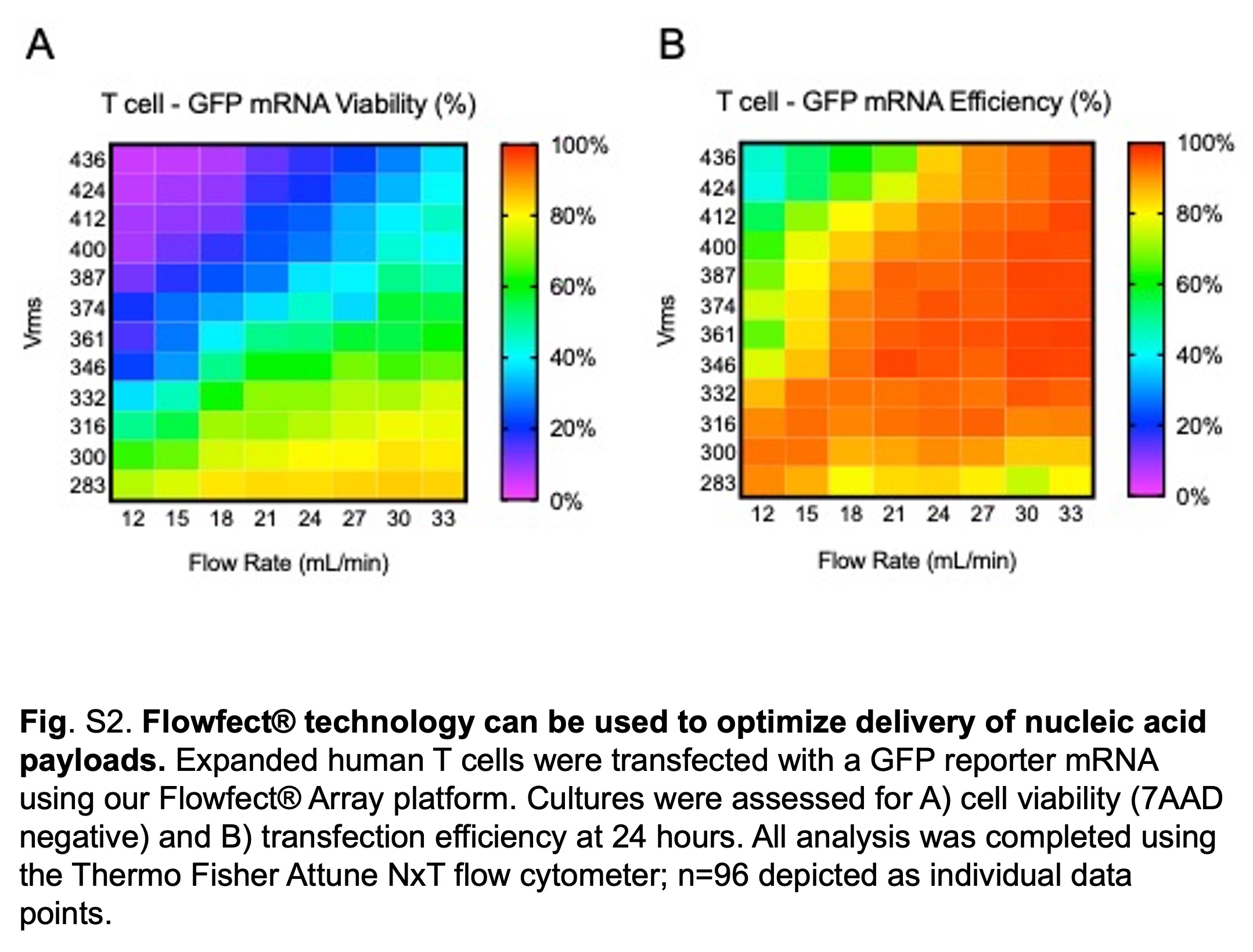

### Supplemental Data 2

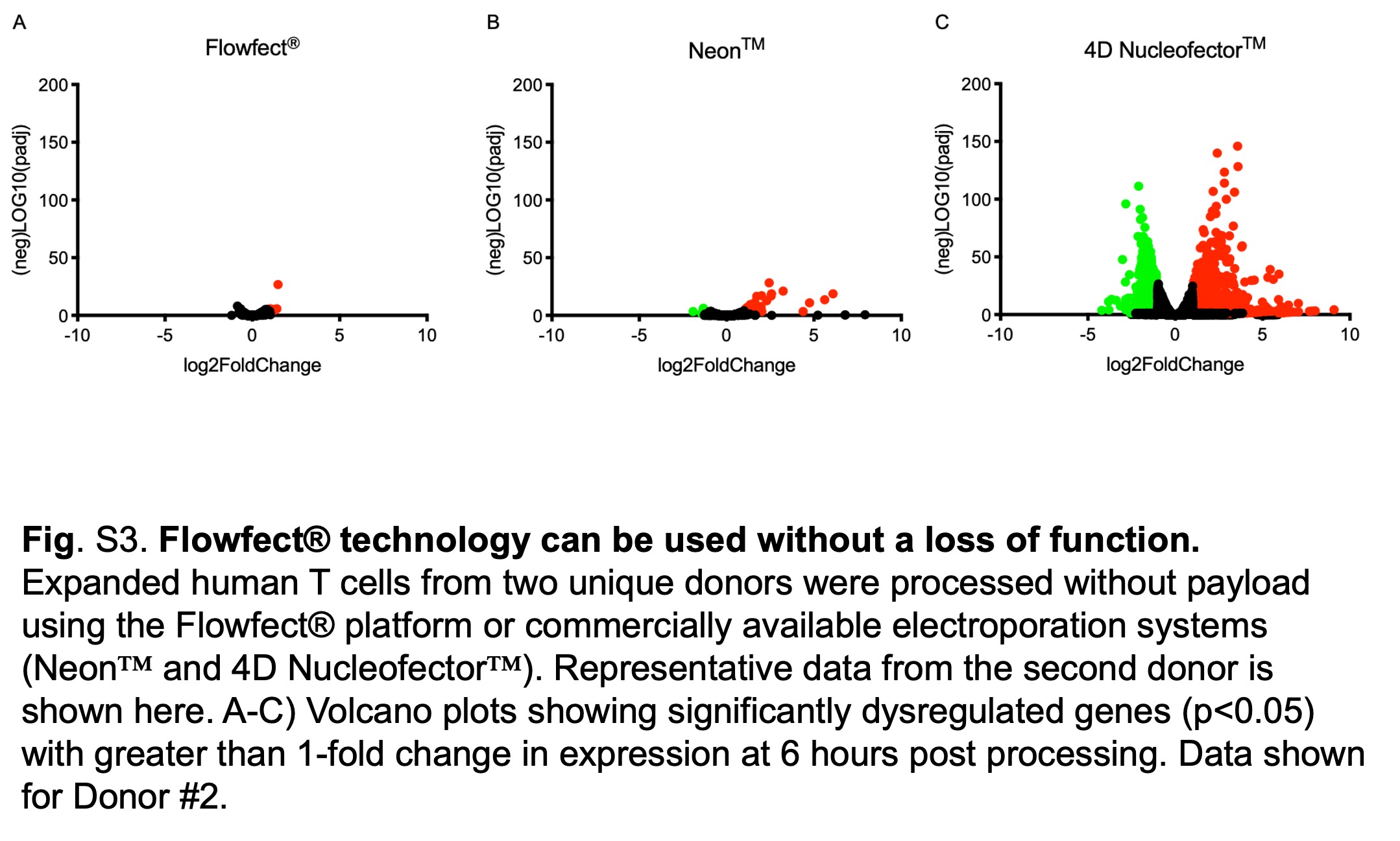

### Supplemental Data 3

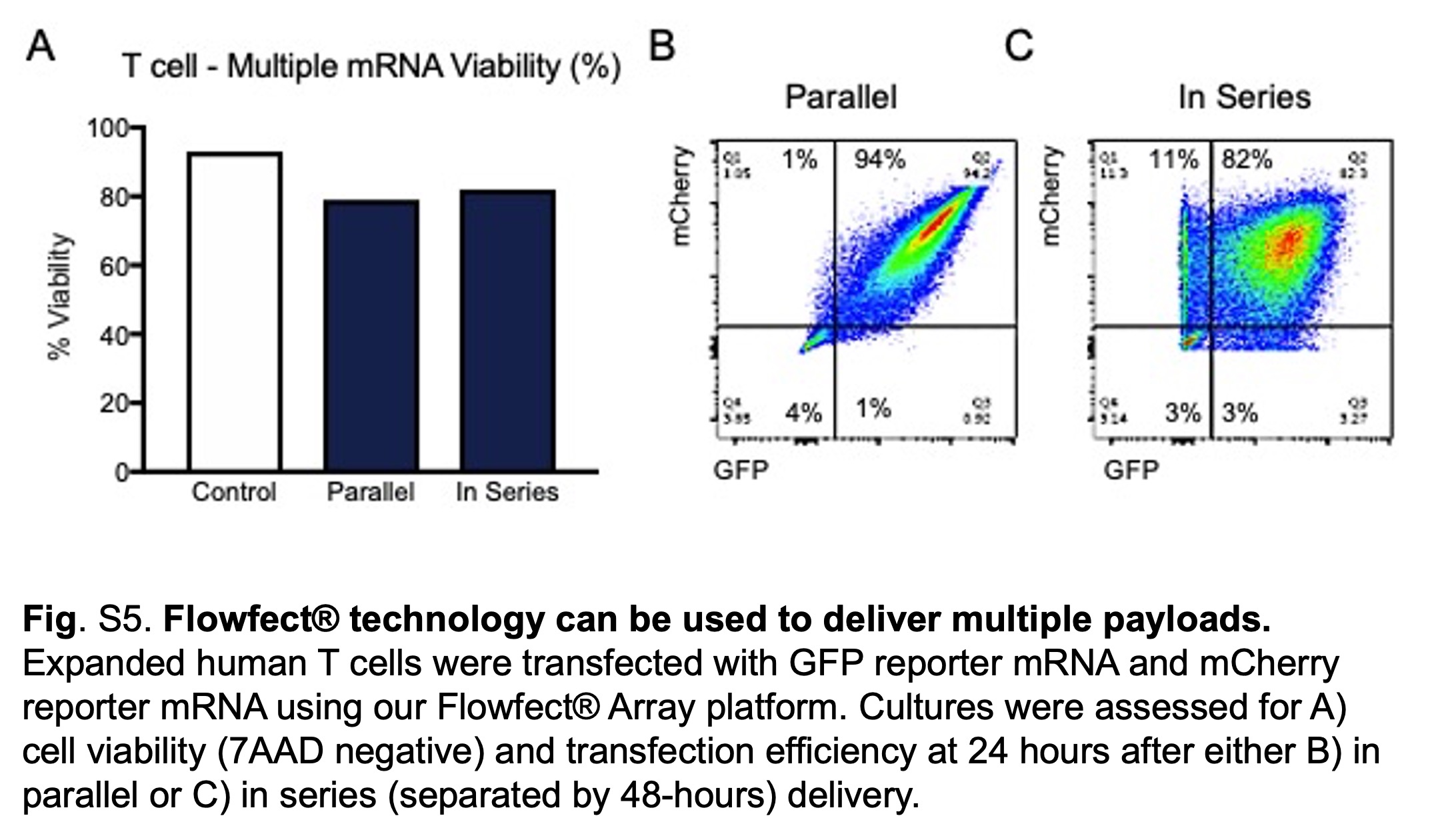

### Table S1) Unique donor demographics supporting data

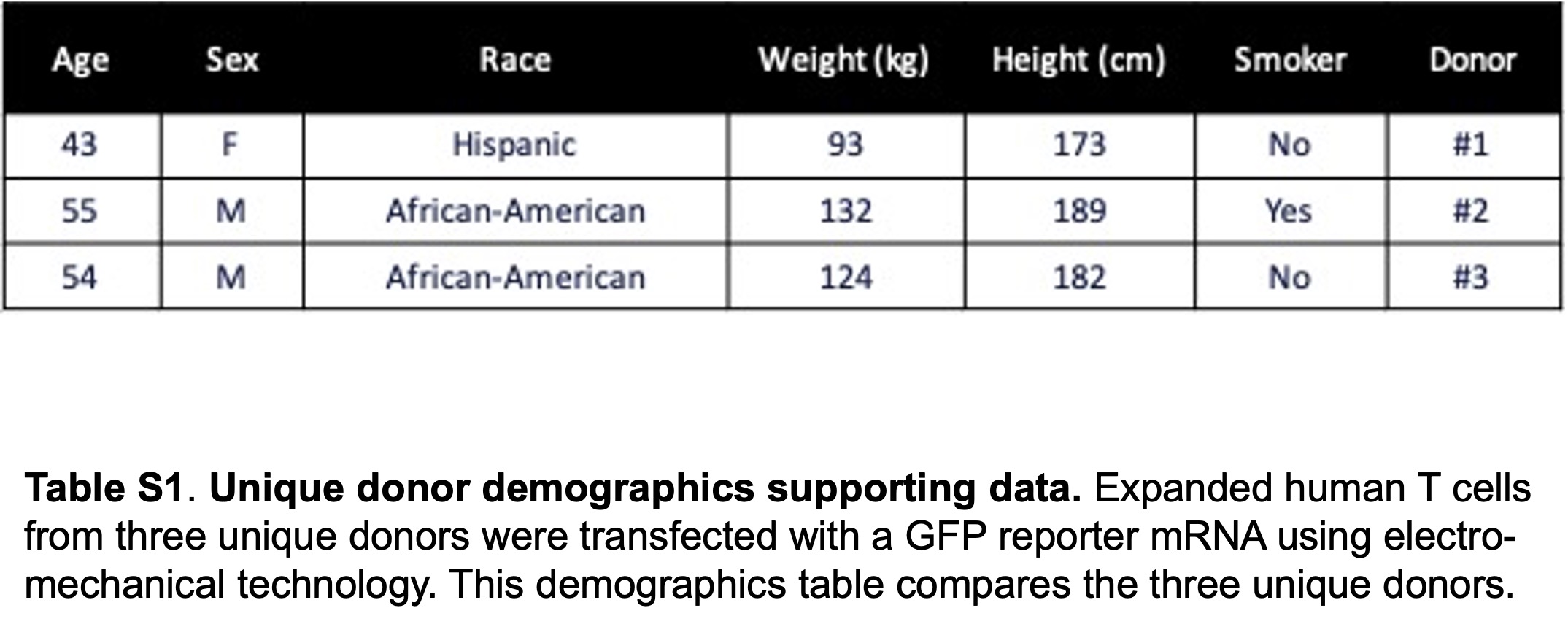
